# A single-cell transcriptome atlas during Cashmere goat hair follicle morphogenesis

**DOI:** 10.1101/2020.01.30.926287

**Authors:** Wei Ge, Wei-Dong Zhang, Yue-Lang Zhang, Yu-Jie Zheng, Fang Li, Shan-He Wang, Jin-Wang Liu, Shao-Jing Tan, Zi-Hui Yan, Lu Wang, Wei Shen, Lei Qu, Xin Wang

## Abstract

Cashmere, also known as soft gold, is produced from the secondary hair follicles in Cashmere goats and it’s therefore of significance to investigate the molecular profiles during Cashmere goat hair follicle development. However, our current understanding of the machinery underlying Cashmere goat hair follicle remains largely unexplored and researches regarding hair follicle development mainly used the mouse as a research model. To provides comprehensively understanding on the cellular heterogeneity and cell lineage cell fate decisions, we performed single-cell RNA sequencing on 19,705 single cells from induction (embryonic day 60), organogenesis (embryonic day 90) and cytodifferentiation (embryonic day 120) stages of fetus Cashmere goat dorsal skin. Unsupervised clustering analysis identified 16 cell clusters and their corresponding cell types were also unprecedentedly characterized. Based on the lineage inference, we revealed detailed molecular landscape along the dermal and epidermal cell lineage developmental pathways. Notably, by cross-species comparasion of single cell data with murine model, we revelaed conserved programs during dermal condensate fate commitment and the heterochrony development of hair follicle development between mouse and Cashmere goat were also discussed here. Our work here delineate unparalleled molecular profiles of different cell populations during Cashmere goat hair follicle morphogenesis and provide a valuable resource for identifying biomarkers during Cashmere goat hair follicle development.

## Introduction

Every year, more than 20,000 tons of Cashmere were generated in China and Cashmere goat has become an important source of income for the people lived in north China (Scott Waldron et al., 2014). Cashmere is the secondary hair follicles in Cashmere goats and forms as early as the fetus stage and due to the commercial value of Cashmere (Ansari-Renani et al., 2011, Geng et al., 2013), it’s therefore of great significance to reveal molecular pathways during early hair follicle development in Cashmere goats. Besides, by using mouse as a research model, research has demonstrated that molecular pathways during early hair follicle morphogenesis play important roles in regulating the hair characteristics, including hair fiber length, fineness and curvature (Duverger & Morasso, 2009), therefore, further enhancing the significance of revealing the molecular pathways driving hair follicle morphogenesis in Cashmere. However, due to the long-time duration of pregnancy (about 145∼159 days) in Cashmere goats (AJ Ritar et al., 1989), researches focused on hair follicle morphogenesis mainly used the mouse as a research model, while our current understanding on hair follicle morphogenesis in Cashmere goat remains largely unknown.

Similar to murine hair follicle development, Cashmere goat hair follicle in uterus development can also be divided into three main stages: induction stage (about embryonic day 55 - 65), organogenesis (about embryonic day 85 - 95) and cytodifferentiation stages (around embryonic day 115) (Zhang Y et al., 2006). In mice, the molecular underpinnings underlying the induction stage has recently been comprehensively investigated by virtue of the development of single-cell RNA sequencing technology, while the late two stages remain not well-known (Saxena et al., 2019). According to what is known in mice, the formation of placodes and dermal condensates (DC) are two marking events in the induction stage and requires a conserved crosstalk between dermal and epidermal cell populations, including Wnt/β-catenin signaling, Edar signaling and Fgf signaling (Chen et al., 2012, Huh et al., 2013, Zhang et al., 2009). More recently, Mok et al., demonstrated that murine DC formation can be further divided into three sub-stages using single-cell RNA sequencing (scRNA seq) technology and characterized detailed transcriptome signature genes during each substage, they also found that ECM/Adhesion signaling was vital for the early DC fate commitment (Mok et al., 2019). The organogenesis stage is characterized by the formation of dermal papilla (DP) cells, hair shaft and inner root sheath (IRS) and the molecular pathways involved includes PDGFα signaling and Shh signaling (Karlsson et al., 1999, Ouspenskaia et al., 2016). For the cytodifferentiation stage, the differentiation of IRS, hair shaft and keratinocyte become obvious and Eda signaling is demonstrated to play a role (Duverger & Morasso, 2009, Millar, 2002). Noteworthy, the asynchronous development of different hair follicles, including guard hair follicles (starts from E13.5), awl, auchene hair follicles (starts from E15.5), and zigzag hair follicle (starts from E17.5) in mice, is also a marking event during the cytodifferentiation stage (Schlake, 2007). However, the molecular machinery underlying the asynchronous development of different hair follicles remains not well-known (Chi et al., 2013, Driskell et al., 2009).

To preliminarily reveal molecular pathways involved during Cashmere goat hair follicle morphogenesis, several groups have collected skin samples from fetus goat and performed transcriptome sequencing analysis to reveal gene expression dynamics between different time points (Gao et al., 2016, Ren et al., 2016). However, due to the lack of conserved markers to label particular cell types within the hair follicles, most studies used skin tissues to perform transcriptome sequencing analysis and generated the “equalized” expression matrices, which is sometimes, hard to reveal the real scenario. The paucity of information regarding the cell heterogeneity within the hair follicles has obviously become the main obstacle in dissecting the hair follicle morphogenesis. scRNA seq has recently became robust tool in dissecting cell heterogeneity and several groups have also successfully used scRNA seq technology to reveal the molecular machinery underlying murine hair follicle development (Ge et al., 2019, Gupta et al., 2019, Mok et al., 2019), further emphasized its application prospect in hair follicle development-related researches.

Tackling the paucity of information regarding the cellular heterogeneity and molecular pathway underlying key cell fate decisions during Cashmere hair follicle development. Here, we reported a single-cell transcriptome landscape during Cashmere goat hair follicle morphogenesis based on 19,705 single-cell transcriptional profiles. We successfully identified different cell types during Cashmere goat hair follicle development and delineated their cell type-specific gene expression profiles, which provides valuable information for the identification of biomarkers and dissecting cellular heterogeneity during Cashmere goat hair follicle development. Besides, cell lineage inference analysis provides a comprehensively understanding of the molecular pathways underlying major cell lineage fate decisions. Our study here provides a valuable resource for understanding Cashmere goat hair follicle development, and will also have implications for future Cashmere goat breeding.

## Results

### Single-cell sequencing and characterization of cellular heterogeneity during Cashmere goat hair follicle morphogenesis

To provide in-depth insight into molecular profiles during Cashmere goat hair follicle development and main cell fates transitions, we collected skin samples from E60, E90 and E120 stage fetus Cashmere goat skin (Supplemental. Fig. S1a), which correspond to hair follicle induction, organogenesis and cytodifferentiation stage and performed single-cell RNA sequencing (Fig. 1a). We totally captured 7,000 single cells for each sample, and for each sample we detected at least 16,000 genes and the genome mapping rate was higher than 90% for all the samples (Supplemental. Fig. S1b). For quality control, we filtered cells according to the number of genes detected (Supplemental. Fig. S1c) and retained high-quality cells for downstream analysis. After quality control, we totally analyzed 19,705 single-cell transcriptome expression profiles from E60 (6,825 single cells), E90 (6,873 single cells) and E120 (6,007 single cells) stage fetus Cashmere goat back skin.

**Fig. 1.**
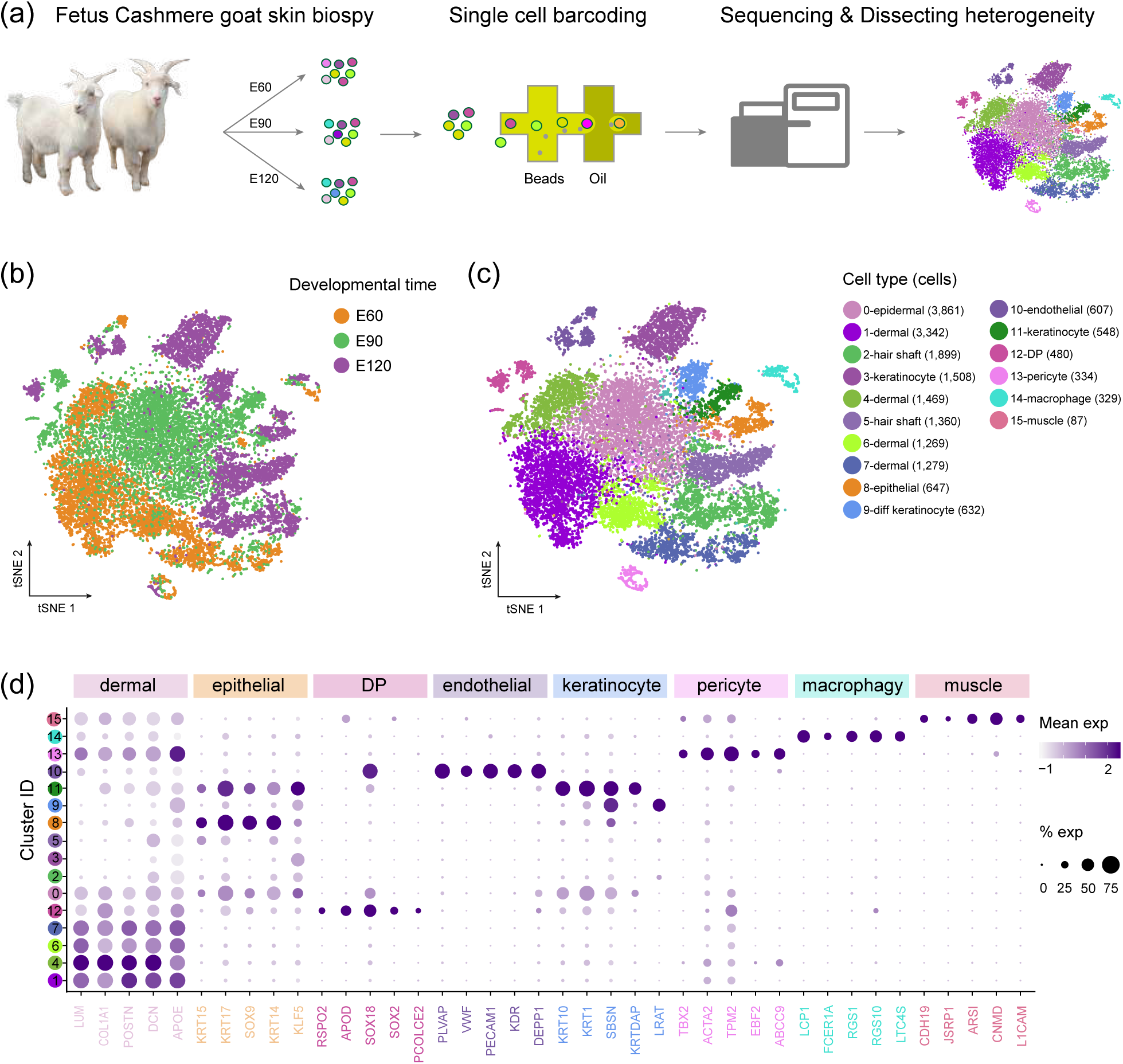
Dissecting cellular heterogeneity during Cashmere goat hair follicle development. (a) Overall experimental design. (b) tSNE plot of all single cells labeled with developmental time. Cells from the different developmental points were color-coded with different colors. (c) tSNE plot of all single cells labeled with cell types according to their marker gene expression. Different colors represent different cell clusters and the cell number for each cluster was listed in the bracket. (d) Dot plot of representative marker genes for different cell clusters. The color intensity represents its expression level, and the dot size represents the positive cell percentage.

To dissect cellular heterogeneity, we next performed t-distributed stochastic neighbor embedding (t-SNE) analysis and we totally identified 16 different cell clusters across three developmental times (Fig. 1b,c and Supplemental. Table 1). By analyzing cluster-specific expressed gene expression, we successfully identified the different cell types according to their marker gene expression (Supplemental. Fig. S1d). Briefly, we found that cluster 1, 4, 6, 7 and 13 expressed high level of dermal cell lineage markers *LUM* and *COL1A1* (Gupta et al., 2019), while cluster 0, 2, 3, 5, 8, 9 and 12 expressed high level of epithelial lineage markers *KRT14* and *KRT17* (Gu & Coulombe, 2007, Joost et al., 2016). Besides, we also identified hair shaft clusters (*LHX2* and *MSX1*, cluster 2, 5) (Yang et al., 2017), endothelial cluster (*KDR* and *PECAM1*, cluster 11) (Detmar et al., 1998), DP cluster (*SOX2* and *SOX18*, cluster 13) (Driskell et al., 2009), pericyte cluster (*ACTA2* and *TPM2*, cluster 15) (Paquet-Fifield et al., 2009), muscle cell cluster (*CNMD* and *ARS1*, cluster 17) and macrophage cell cluster (*ALF1* and *RGS1*, cluster 16) (Lee et al., 2016). More importantly, we further delineated transcriptional characteristics for each cell types and identified a series of cell type-specific marker genes during Cashmere goat hair follicle development (Fig. 1d), and it was also worth noting that many cell type-specific expressed marker genes were also consistent with a murine scenario, such as dermal cell markers *POSTN, DCN, APOE*, epithelial cell markers *KRT14, KRT15*, and DP cell markers *SOX2, SOX18*.

### Defining dermal cell lineage and epidermal cell lineage developmental trajectory along pseudoptime

After the characterization of different cell clusters, we then want to investigate major cell fate transitions during hair follicle development. We, therefore, performed pseudotime trajectory construction analysis on dermal and epidermal cell clusters (Fig. 2). Since we have successfully characterized all cell clusters, we then selected dermal cell lineage cell clusters (Fig. 2a, cluster 1, 4, 6, 7 and 12) and epidermal cell lineage cell clusters (Fig. 2b, cluster 0, 2, 3, 5, 8, 9 and 11) to infer cell lineage developmental trajectory. For the dermal cell lineage, pseudotime trajectory displayed 2 branch points (Fig. 2c), while the epidermal cell lineage showed 3 branch points (Fig. 2d). Noteworthy, when the cells were color-coded with their corresponding developmental time, they also showed a time-ordered pattern along the pseudotime. As for the branch point in the dermal and epidermal cell populations, based on the prior knowledge on cell lineage dynamics, during early hair follicle development, the dermal cell fate involves DC fate commitment and DP fate commitment (Fig. 2e) (Saxena et al., 2019), while epidermal cell fate involves matrix cell fate commitment, hair shaft/IRS fate commitment and keratinocyte fate commitment (Fig. 2f) (Forni et al., 2012, Millar, 2002, Schmidt-Ullrich & Paus, 2005), it’s therefore different branch points may represent the process of cell fate decisions.

**Fig. 2.**
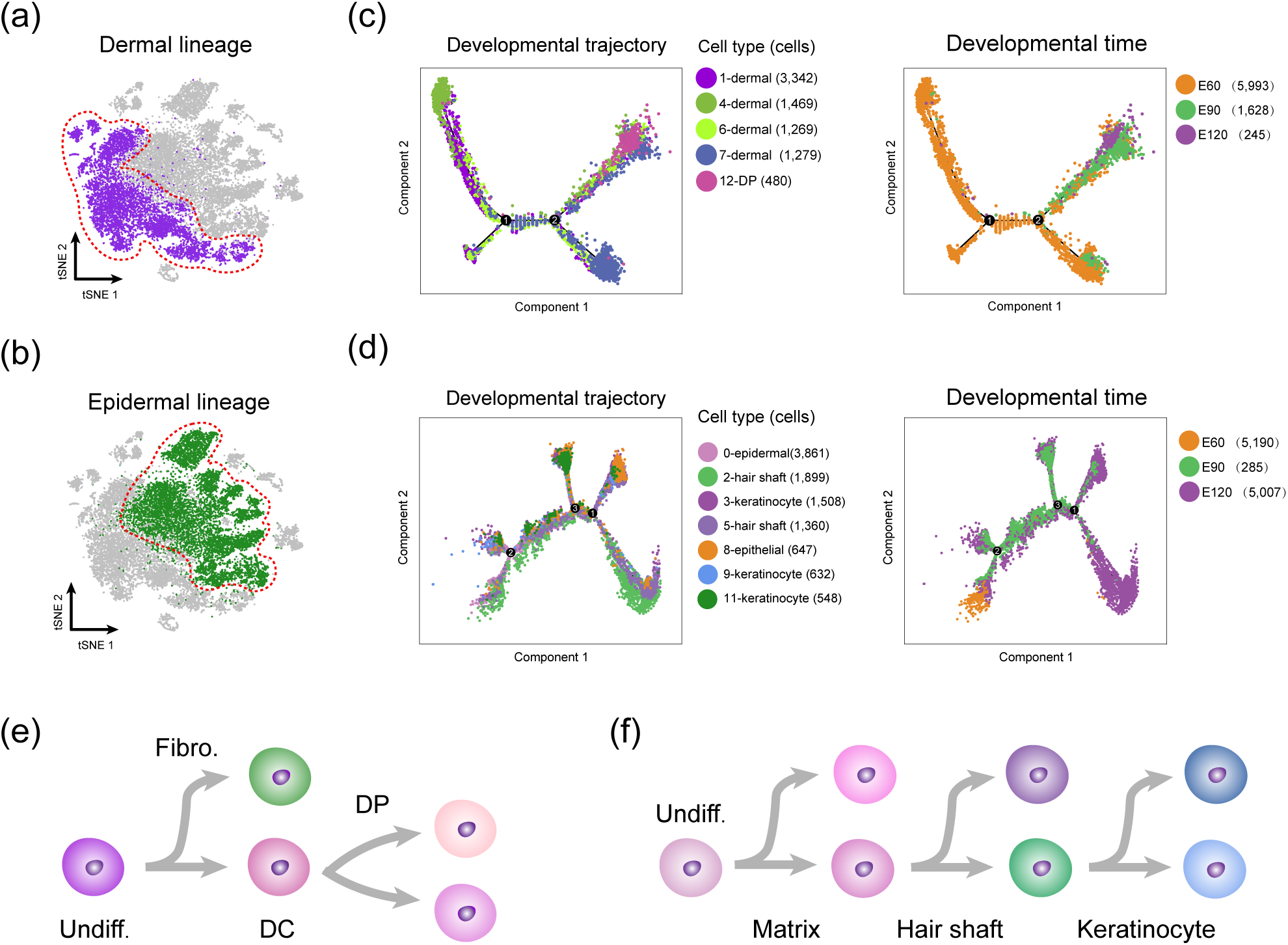
Delineating dermal and epidermal cell lineage pseudotime developmental trajectory. (a) Dermal cell lineage highlighted in the tSNE plot. (b) Developmental trajectory of dermal cell lineage along pseudotime. Cells were color-coded with cell types identified by Seurat (left panel) and developmental time (right panel), respectively. (c) Epidermal cell lineage highlighted in the tSNE plot. (d) Developmental trajectory of epidermal cell lineage along pseudotime. Cells were also color-coded with cell types (left panel) and developmental time (right panel), respectively. (e) Main dermal cell lineage decisions during hair follicle morphogenesis. (f) Main epidermal cell lineage fate decisions during hair follicle morphogenesis.

### Delineating developmental pathway during DC fate commitment

After dermal cell lineage trajectory inference, we firstly focused on the first branch point on the dermal cell pseudotime trajectory to reveal the first dermal cell fate decision. By analyzing gene expression dynamics along pseudotime, we observed 2,679 differentially expressed genes at the end of cell fate commitment (Supplemental. Table 2) and gene functional enrichment analysis revealed that these genes enriched GO terms of “tissue morphogenesis, response to growth factor and cell morphogenesis involved in differentiation” (Fig. 3a). A comparison of overlapped GO terms between each gene set showed that they shared substantial GO terms (Supplemental. Fig. 2). Of particular notice, we observed a series of canonical murine dermal condensate cell markers such as *APOD, LUM* and *APOE* (Mok et al., 2019) in those differentially expressed genes, and the pseudotime expression pattern of DC markers *APOD, SOX18, CTNNB1* and *SOX2* was increased along pseudotime (Fig. 3b), therefore, we termed the first branch point as DC fate commitment.

**Fig. 3.**
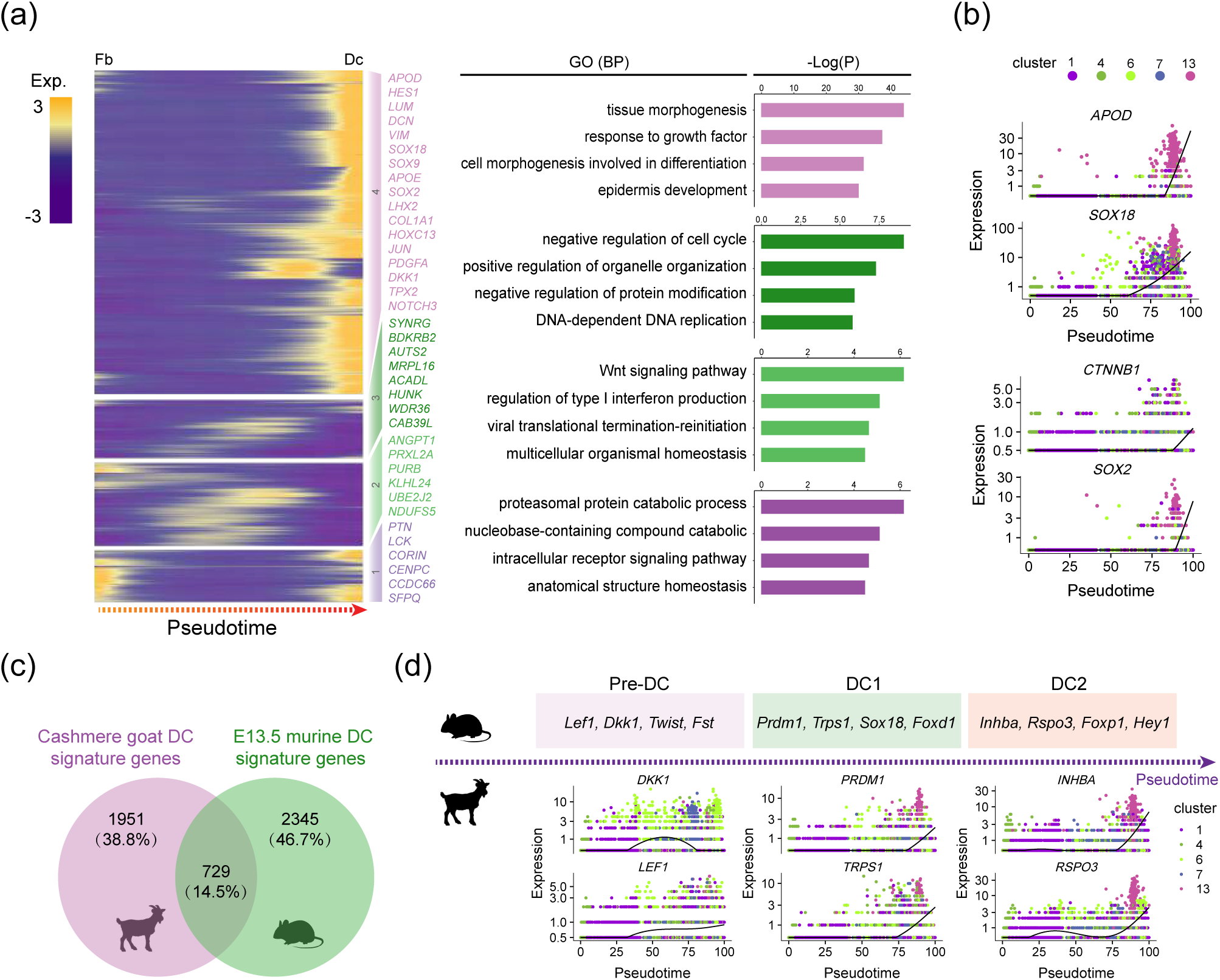
Revealing molecular profiles during DC fate commitment. (a) Pseudotime expression heatmap during DC fate commitment. The four gene sets were determined by k-means clustering according to their expression pattern and GO terms for each gene set were listed in the right panel. (b) Immunofluorescence analysis of BMP2, CTNNB1, LEF1 and K15 in E60 Cashmere goat skin tissues. Scale bars = 50 μm. (c) Veen diagram illustrating overlapped signature genes between E60 Cashmere goat DC cells and murine E13.5 DC signature genes. (d) Visualizing murine DC signature genes of different stages along pseudotime in Cashmere goat. The murine signature DC markers were listed in the top panel and their corresponding expression pattern in Cashmere goat was listed in the lower panel. Cells were color-coded according to their cluster identify.

To our knowledge, no reports yet have delineated cellular heterogeneity and transcriptional landscape during Cashmere goat hair follicle morphogenesis, and most researches regarding hair follicle development have been performed on the murine model. To gain in-depth insight into machinery driving DC fate commitment, we then compared DC fate signature genes with recently reported murine DC fate commitment signature genes (Fig. 3c) and observed 729 (about 14.5% of all) overlapped genes between Cashmere goat and murine DC signature genes (Supplemental. Table 3). Noteworthy, based on single-cell RNA sequencing on E15.0 dorsal skin, Mok et al. recently demonstrated that murine DC fate commitment can be divided into pre-DC, DC1 and DC2 stage at a more detailed level (Mok et al., 2019). By analyzing murine DC marker genes at different stages, we similarly found that DC signature genes of Cashmere goat also showed chronological expression patterns along pseudotime (Fig. 3d). Briefly, murine pre-DC markers *DKK1* and *LEF1* were elevated prior to DC1 and DC2 marker expressions, such as *PRDM1, TRPS1, INHBA* and *RSPO3*, it’s therefore plausible that DC fate commitment in Cashmere goat may also involve different stages.

### Delineating DP cell heterogeneity along pseudotime

After defining DC fate commitment, we then focused on the next branch point. We firstly compared differentially expressed gene expression between the two branches and observed that cell fate 1 elevated canonical DP marker genes, such as *APOD, SOX18*, and enriched GO terms of “tissue morphogenesis and epidermis development”, while cell fate 2 elevated genes such as *CENPW, TOP2A* and enriched GO terms of “mitotic cell cycle process and DNA-dependent DNA replication” (Fig. 4a,b and Supplemental. Fig. 3a,b). To gain an in-depth understanding of the differences between the two branches, we further identified differentially expressed genes between the two branches using another differential analysis method (Wilcox, Wilcoxon Rank Sum test). Consistent with Monocle analysis, cell fate 1 also showed higher expression of *APOD, SOX18* and *IGF1*, while cell fate 2 showed elevated expression of *CENPW, TOP2A* and *DCN* (Fig. 4c). GO enrichment analysis similarly revealed that differentially expressed genes in cell fate 1 enriched GO terms of “tissue morphogenesis and epithelial cell differentiation”, while cell fate 2 enriched GO mainly related to the regulation of cell cycle (Supplemental. Fig. 3c).

**Fig 4.**
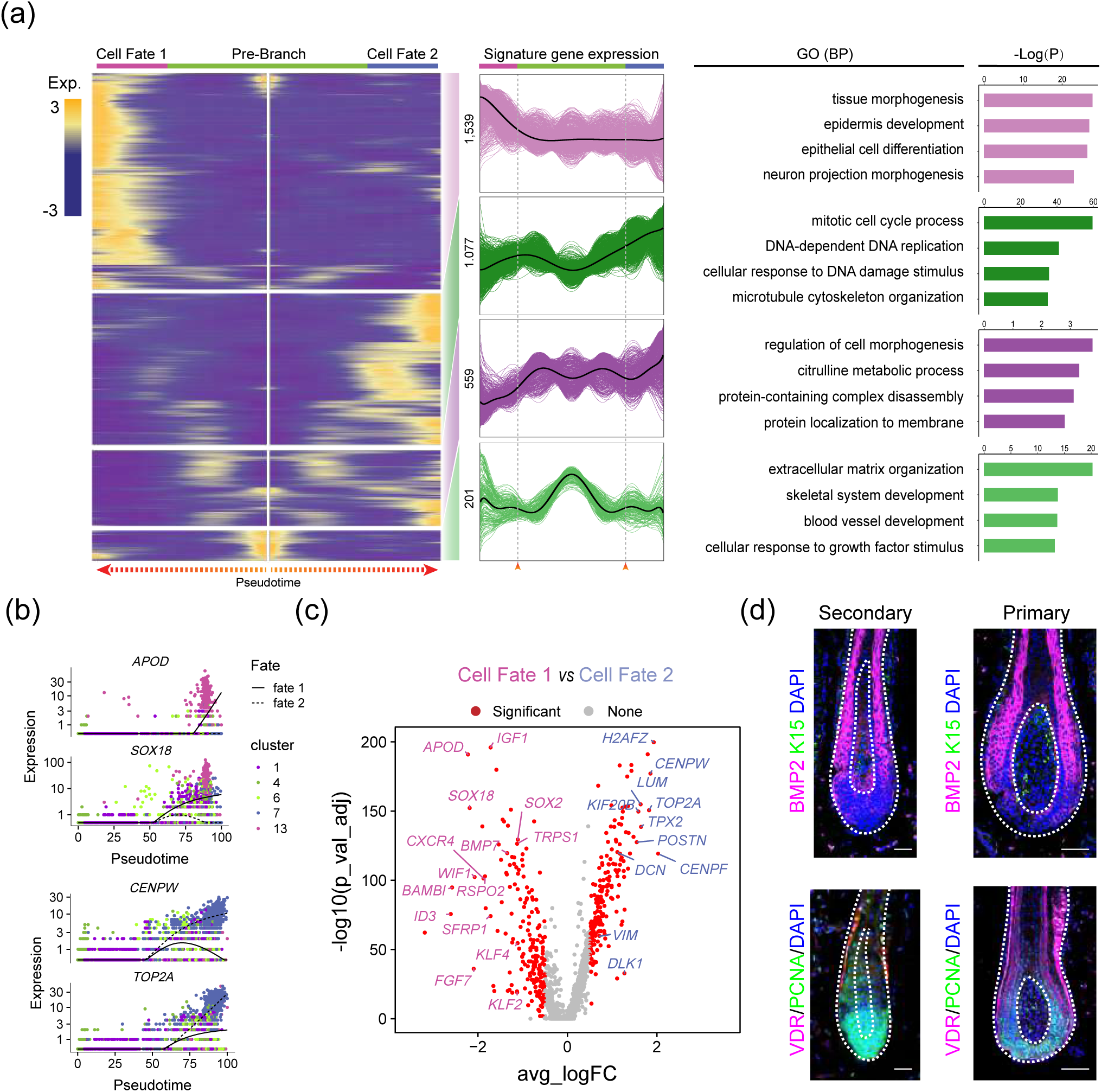
DP cells from primary hair follicles and secondary hair follicles in Cashmere goat showed distinct gene expression profiles. (a) Heatmap illustrating dynamic gene expression profiles during DP cell fate commitment. The gene expression pattern for each gene set was listed in the middle panel, and the top 5 enriched GO terms for each gene set were listed in the right panel. (b) Pseudotime expression of cell fate 1 enriched genes *APOD, SOX18* and cell fate 2 enriched genes *CENPW, TOP2A*. Cells were colored-coded with their corresponding cluster identity. (c) Volcano plot illustrating cell fate 1 and cell fate significantly enriched differentially expressed genes. (d) Immunofluorescence analysis of BMP2, K15, VDR and PCNA in the primary hair follicles and secondary hair follicles from E120 Cashmere skin sections. Scale bars = 50 μm.

To dissect the heterogeneity within DP cells, we further performed immunofluorescence analysis on E120 Cashmere goat back skin tissues and observed different staining patterns between primary hair follicles (PF) and secondary hair follicles (SF) (Fig. 4d). For the BMP2 staining, we found that BMP2 specifically expressed in the matrix cells surrounding DP cells, while in the secondary hair follicle, BMP2 positive cells could be found in both DP cells and surrounding matrix cells. Besides, we found that PCNA, a cell cycle-related marker, was expressed mainly in surrounding matrix cells in the primary hair follicle, while in the secondary hair follicles, we observed high expression of PCNA both in DP cells and surrounding matrix cells. Taken together, our analysis here demonstrated that different hair follicles showed different gene expression patterns in Cashmere goat, which may also plausible for the asynchronous development of different hair follicles.

### Delineating developmental pathway during the first epidermal cell fate decision

After revealing dermal cell fate decisions, we then focused on the epidermal cell clusters. Monocle pseudotime trajectory inference analysis revealed that epidermal cells showed three different branch points. We then firstly focused on the first branch point and analyzed differentially expressed gene dynamics along pseudotime. Based on k-means clustering, we observed four distinct gene clusters. As expected, we observed a series matrix cell markers at the end of pseodotime, including *HOXC13, KRT25* and *KRT71*, thus deciphering a matrix cell fate commitment (Fig. 5a). Gene functional enrichment analysis showed that matrix cell expressed signature genes enriched GO terms of “keratinocyte differentiation, epidermis development, and skin epidermis development” (Fig. 5b), while comparison of GO terms between different gene set reveals that gene set 1,2 differs to that of gene set 3,4 (Supplemental. Fig. 4b).

**Fig. 5.**
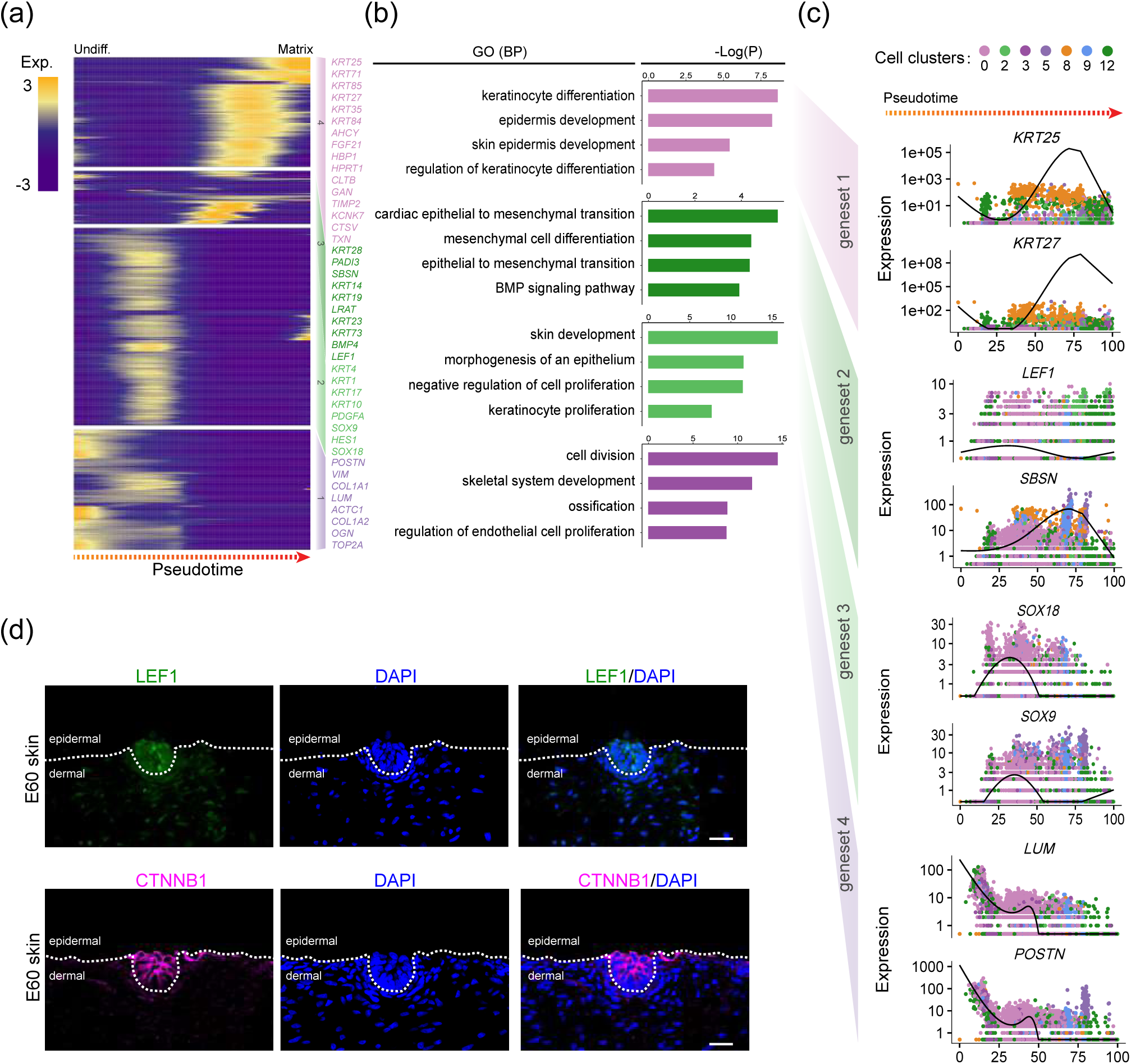
Delineating matrix cell fate decision along pseudotime. (a) Heatmap demonstrating dynamic gene expression patterns during matrix cell fate commitment. The differentially expressed genes were divided into 4 different gene sets based on k-means clustering. (b) Gene functional enrichment analysis of signature genes in the 4 different gene sets identified in Fig. 5a. (c) The pseudotime expression pattern of representative signature genes from each gene set. Cells were color-coded according to their cluster identity. (d) Immunofluorescence analysis of LEF1 and CTNNB1 in E60 Cashmere skin tissues. Scale bars = 50 μm.

To gain in-depth insight into molecular profiles during matrix cell fate commitment, we further analyzed the signature gene expression pattern along pseudotime. For the gene set 4, namely matrix cell fate, they enriched a series of keratin family genes, such as *KRT25, KRT27, KRT84*, and their expression was increased along pseudotime (Fig. 5c). For the gene set 3 and 2, we found that they transiently elevated expression of *LEF1, SBSN, SOX18, SOX9*, and enriched GO terms of “cardiac epithelial to mesenchymal transition, mesenchymal cell differentiation and skin development, morphogenesis of an epithelium, respectively. For the gene set 1, they enriched genes such as *VIM, LUM* and *COL1A1* and both showed decreased expression along pseudotime. Immunohistochemistry staining analysis also confirmed that LEF1 and CTNNB1 were expressed in the upper epidermis, which was also consistent with a murine scenario (Polakis, 2001, Tsai et al., 2014).

### Delineating cell fate decisions during hair shaft and IRS cell fate commitment

After deciphering matrix cell fate commitment in the first epidermal trajectory point, we next focused on the next branch point. By analyzing differentially expressed genes along pseudotime, we found that cell fate 1 enriched canonical hair shaft markers, including *SHH, VDR*, and *HOXC13*, while cell fate 2 enriched canonical IRS markers, such as *SOX9, KRT14, SBSN* and *LEF1* (Fig. 6a) (Yang et al., 2017). Our immunofluorescence results also confirmed their expression in the Cashmere hair follicles (Supplemental. Fig. 5a). For the pre-branch, we observed genes such as *LUM, COL1A, OGN, SOX18* and they all showed decreased expression along pseudotime (Fig. 6b). For the hair shaft fate (cell fate 1), we also observed elevated expression of *DCN, TOP2A, CENPW* and *H2AFZ* and enriched GO terms of “mitotic cell cycle process and DNA replication” (Supplemental. Fig. 5 a,b). For the IRS cell fate (cell fate 2), we observed elevated expression of *PRDM1, KRT1, SOX9, KRT14* and enriched GO terms of “supramolecular fiber organization and tissues morphogenesis”.

**Fig. 6.**
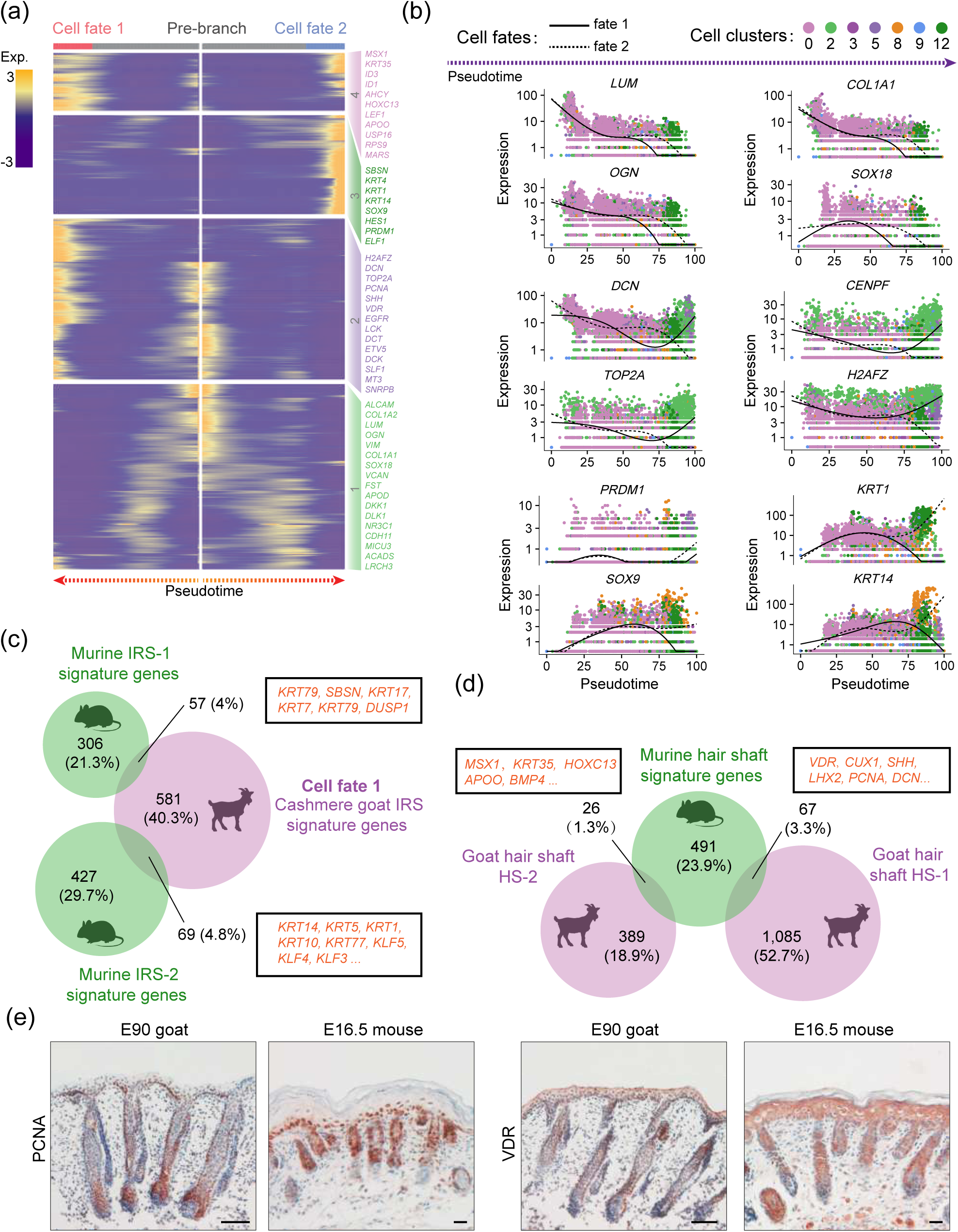
Revealing pseudotime gene expression dynamics during IRS and hair shaft cell fate commitment. (a) Heatmap illustrating pseudotime gene expression pattern of differentially expressed genes during IRS and hair shaft cell development. Cell fate 1 depicts IRS fate, while cell fate 2 depicts hair shaft fates. (b) Pseudotime expression pattern of representative marker genes during IRS and hair shaft cell fate commitment. Cells were color-coded with their corresponding cluster identity and the solid line depicts cell fate 1, while the dashed line depicts cell fate 2. (c) Veen diagram demonstrating overlapped murine and Cashmere goat IRS signature genes. The representative overlapped genes were listed in the corresponding boxes. (d) Comparison of murine hair shaft signature genes with Cashmere goat hair shaft signature genes. The representative overlapped genes were listed in the rectangular boxes. (e) Comparison of PCNA and VDR expression in E90 Cashmere goat and E16.5 mouse skin tissues. Scale bars = 50 μm.

Besides, we also compared our identified hair shaft and IRS signature genes with our recently identified murine hair shaft and IRS signature genes (Ge et al., 2019). Interestingly, we only observed about ∼8.8% overlapped IRS signature genes between mouse and goat (Fig. 6c and Supplemental. Table 4), and these overlapped genes mainly consisted of Keratin family genes, such as *KRT14, KRT17, KRT79*. As for overlapped hair shaft signature genes, we only observed ∼4.6% genes (Fig. 6d), including *MSX1, HOXC13, APOD, LHX2, SHH*, and *VDR*. To further validate our analysis, we compared the expression of PCNA and VDR between E90 Cashmere goat skin and E16.5 mouse skin tissues, and the immunohistochemistry showed that they showed similar expression patterns during hair follicle development (Fig. 6e). These data together demonstrate that hair shaft and IRS specification may require a conserved program during cell fate decisions.

### Revealing the developmental pathway during keratinocyte cell fate commitment

After delineating the first two epidermal cell lineage cell fate decisions, we then focused on the last branch point. Pseudotime trajectory analysis also revealed two different branches, and we then analyzed pseudotime gene expression dynamics between the two branches (Fig. 7a). Analyzing differentially expressed genes along psusotime, we found that cell fate 1 enriched genes such as *VDR, BMP4, STAR, KRT85* and *KRT14*, while cell fate 2 enriched genes such as *BMP2, SHH, CUX1* and *ETV5*. Noteworthy, *KRT14* has been identified as a marker for keratinocyte both in humans and mice (Green et al., 2003, Joost et al., 2016), we, therefore, termed this fate as keratinocytes. To gain in-depth insight into their corresponding cell type, we performed gene functional enrichment analysis for each gene set and analyzed pseudotime GO enrichment dynamics (Fig. 7b and Supplemental. Fig. 6a). The result showed that the pre-branch mainly enriched genes related to “regulation of response process”, while for the cell fate 1, the signature genes mainly enriched GO terms of “development differentiation of epidermal”. Interestingly, for cell fate 2, these differentially expressed genes mainly enriched in GO terms of “regulation of cell cycle” and showed lower expression of *KRT14* (Fig. 7c), it’s therefore plausible that keratinocyte differentiation at this stage is not synchronized. To confirm such hypothesis, we performed immunofluorescence analysis of VDR, PCNA, BMP2, CTNNB1 and LEF1, and the results showed that the expression of VDR and BMP2 in the outer layer epidermis was not homogeneous, with some cell clusters showed higher expression while some cell populations showed lower expression (Fig. 7d). For the pre-branch enriched CTNNB1, it was uniformly expressed in the interfollicular epidermis. Besides, the expression pattern of PCNA and LEF1 was even more significant and they were partially expressed in the epidermis. Taken together, these results together emphasize that keratinocyte differentiation in Cashmere goat is asynchronous and requires different gene expression profiles.

**Fig. 7.**
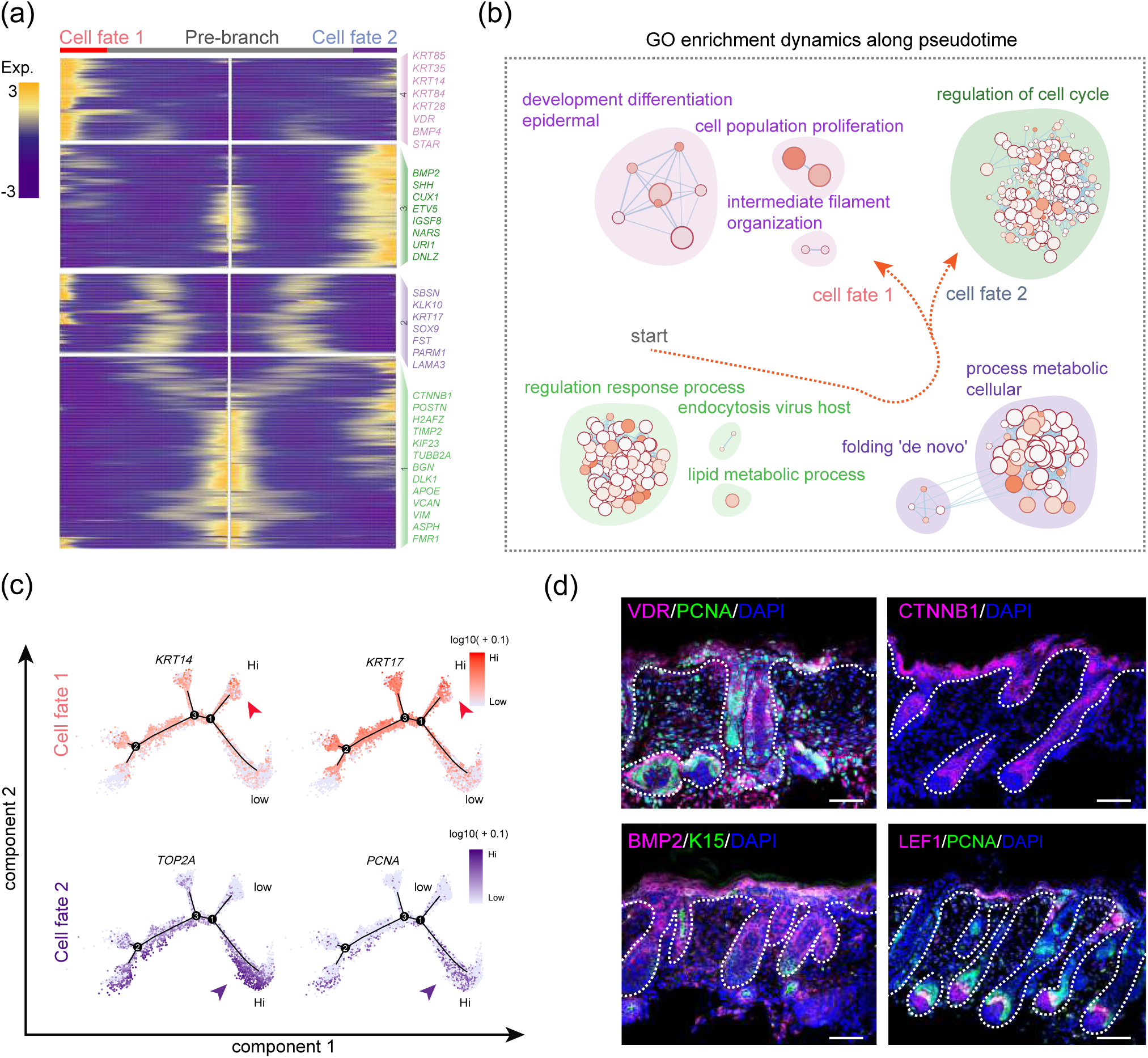
Dissecting the asynchronous development of keratinocyte. (a) Heatmap illustrating gene expression dynamics during keratinocyte differentiation. (b) GO terms corresponding to the gene set in Figure 7a. The arrow indicates the elongation of pseudotime. (c) Expression of cell fate 1 signature genes *KRT14, KRT17* and cell fate 2 signature genes *TOP2A, PCNA* along the pseudotime trajectory. (d) Immunofluorescent staining analysis of VDR, PCNA, CTNNB1, BMP2, LEF1 and K15 in E90 Cashmere goat skin tissues. Scale bars = 50 μm.

## Discussion

The development of scRNA seq in recent years has been demonstrated as a robust tool for the developmental biologist to investigate organogenesis and has provided us with unparalleled insight into mammalian development. In the past decades, the number of papers using scRNA seq based technology has increased exponentially and scRNA has also been awarded Science’s 2018 Breakthrough of the Year (Angerer et al., 2017, Pennisi, 2018). Here, we successfully constructed single-cell atlas during Cashmere goat hair follicle development. Based on the downstream analysis, we provided unparalleled insight into the cellular heterogeneity and major cell fate decisions during the Cashmere hair follicle in uterus morphogenesis. As far as we have known, this is the first study to comprehensively delineating molecular profiles of various cell types and revealing major cell fate decisions during Cashmere hair follicle development. More importantly, by analyzing cluster-specific gene expression profiles, our data here provides a valuable resource for identification of markers for future studies in Cashmere skin tissues and also provides an in-depth understanding of the Cashmere hair follicle development.

In the current study, we totally analyzed 19,705 single-cell transcriptional profiles from three different points, which can represent major cell types during hair follicle development. To dissect cellular heterogeneity during Cashmere goat hair follicle development after tSNE analysis, we analyzed cluster-specific expressed signature genes for each cluster to infer their corresponding cell types. It’s worth noting that all the cell markers used in the current study were referenced from the murine model due to the paucity of information regarding marker gene expression during Cashmere goat hair follicle development. However, our study here showed that substantial murine cell type-specific biomarkers are identical to the Cashmere goat. Besides, by comparison of cell type-specific signature genes between Cashmere goat and mouse (Fig. 3c and Fig. 6c,d), we found that the transcriptome similarity decreased along pseudotime (revealed by overlapped signature genes in this study). Briefly, about 729 DC signature genes overlapped between Cashmere goat and mouse, and murine DC signature genes at different stages (pre-DC, DC1 and DC2) showed similar expression pattern along pseudotime in Cashmere goat (Mok et al., 2019). While for the late-stage IRS and hair shaft cell population in the late stage of hair follicle development, the percentage of overlapped genes decreased (126 and 93 overlapped genes, respectively). Similarly, by comparing gene expression across 7 different species from brain, heart, ovary, kidney, testis and liver, researches have demonstrated that organ-specific molecular profiles are more similar in early development and become more distinct during development (Cardoso-Moreira et al., 2019), it’s therefore plausible that hair follicle development in Cashmere goat and mouse may also consistent with such theory.

Similar to the murine model, we also observed differential gene expression profiles in different DP cell populations from different hair follicles (primary hair follicles vs secondary hair follicles), which was also consistent with our recent findings on mice (Ge et al., 2019). Besides, an in-depth comparison of differentially expressed genes revealed that different DP cell populations require different transcriptional profiles, and it’s, therefore, plausible that the different induction signals may be involved during the asynchronous development of hair follicles. Consistent with such hypothesis, researches found that SOX2 was specifically expressed in guard hair follicles, but not zigzag hair follicles, while *SOX18* was demonstrated to regulate zigzag hair follicle morphogenesis (Driskell et al., 2009, James et al., 2003, Pennisi et al., 2000). Although our study here also demonstrates the asynchronous development of different hair follicles in Cashmere goat, the detailed machinery underlying such phenomenon was not investigated here and future studies may focus on such topics, which will definitely provide us new insight into the hair follicle biology and hair follicle regeneration.

Based on single-cell pseudotime trajectory inference, our study here also highlighted the underappreciated cell fate decisions during hair follicle development. Different from our previous studies performed in mice, our pseudotime lineage trajectory of epidermal cell lineage showed three different branch points while murine epidermal cell lineage trajectory showed two branch points. by analyzing the gene expression profiles of the first two branch points, they actually showed similar cell fate decisions, while the additional branch point in Cashmere goat mainly comes from keratinocyte differentiation. Such differences in pseudotime trajectory may be partially explained by the differences in the development of hair follicles across species, for example, E120 stage Cashmere goat showed obvious hair fiber on the surface of the skin, while it was not until 6-7 postnatal day that the hair follicles in the mouse skin surface become visible. The difference in the pseudotime trajectory reveals that hair follicle development was heterochrony between Cashmere goat and mouse, which was frequent when comparing the development of specific organs across species.

Finally, our dataset here provides an important resource for understanding the cellular heterogeneity and major cell fate decisions during Cashmere goat hair follicle development. For the first time, the detailed transcriptional landscape of different cell populations was delineated here at single-cell resolution, which provided valuable resources for the identification of biomarkers. Besides, our trajectory inference analysis here successfully recapitulated major cell fate decisions during Cashmere goat hair follicle development, which enabled us to comprehensively study the detailed developmental pathways involved during Cashmere goat hair follicle morphogenesis, and will also have implications for the Cashmere goat breeding work in animal husbandry.

## Materials and methods

### Experimental Animals

All the experimental Shaanbei White Cashmere goats involved in this study were obtained from the Shaanbei Cashmere Goat Engineering Technology Research Center of Shaanxi Province and were fed with Cashmere goat feeding standard (DB61/T583-2013) of Shaanxi Province. All pregnant goats were prepared using artificial insemination and all the experimental procedures involved goats in this study were approved by the Experimental Animal Manage Committee of Northwest A&F University.

### Single-cell suspension preparation

The goat fetus at desired dates was isolated using cesarean operation when the pregnant goats were anesthetic with the compound ketamine. The skin tissues (0.5 cm × 0.5 cm) were isolated from the fetus back skin and were immediately transferred to the ice-cold DMEM/F12 media (Gibco, Beijing, China) with 50 U/ml penicillin and 50 mg/ml streptomycin (HyClone, Beijing, China). After washing three times with DMEM/F12 to remove contaminative blood cells, the skin tissues were then dissociated into single cells prior to sequencing. For E60 and E90 skin tissues, the obtained skin tissues were firstly incubated with 2 mg/ml collagenase IV (Sigma, St Louis, MO, USA) for 30 min at 37°C, and then the skin tissues were mechanically dissociated into single-cell suspensions with a 1ml pipette tip. For E120 skin tissues, the skin tissues were firstly cut into ∼3 mm skin pieces and then incubated with 2 mg/ml collagenase IV for 30 min. After incubation, the hair follicles within the skin tissues were isolated with a pair of precise forceps and the pooled hair follicles were further dissociated into single cells with TypLE Express (Gibco, Grand Island, NY, USA) for 30 min at 37°C. The obtained single-cell suspensions were then washed three times with PBS supplemented with 0.04% BSA (Sigma, St Louis, MO, USA) and were filtered with a 40 μm cell strainer (BD Falcon, BD Biosciences, San Jose, CA, USA) to remove debris and cell aggregations. For each stage, the samples were obtained from at least 2 different goat fetus and for each fetus goat, the single-cell suspension was prepared separately until finally pooled together prior to single-cell barcoding.

### Single-cell library construction and sequencing

Single-cell library was constructed using 10x Genomics’ Chromium Single Cell 3’ V3 Gel Beads Kit (10x Genomics, Pleasanton, CA, USA) according to the manufacture’s instructions. After cell counting, the single-cell suspension was adjusted to 1000 cells/μl and about 7,000 cells were obtained for each stage. The single-cell barcoding procedure was performed using a 10x Genomics Chromium barcoding system (10x Genomics, Pleasanton, CA, USA) according to the manufacturer’s guide. After single-cell library construction, the Illumina HiSeq X Ten sequencer (Illumina, San Diego, CA, USA) was used to sequence and 150 bp pair ended reads were generated.

### 10x Genomics single-cell RNA sequencing data processing

The obtained raw sequencing files were processed with standard CellRanger (v2.2.0) pipeline according to the 10x Genomics official guide (https://www.10xgenomics.com/cn/). The produced raw base call files were firstly transformed into fastq files by using Cellranger mkfastq function. The goat ARS1 reference genome downloaded from ensemble was used as a reference genome (https://asia.ensembl.org/Capra_hircus/Info/Index). Cellranger count function was used to perform mapping, filtering low-quality cells, barcoding counting and UMI counting.

After the standard Cellranger pipeline, the generated gene expression matrice files were then analyzed with Seurat (V2.3.4) package according to the official user guide (https://satijalab.org/seurat/vignettes.html). Quality control was performed using FilterCells function and cells with detected genes less than 200 and genes expressed less than 3 cells were filtered. After normalization and data scaling, the different datasets were integrated by using RunMultiCCA function. tSNE was used to perform dimension reduction analysis and different cell clusters were identified by using FindClusters function. The cluster specifically expressed genes were analyzed with FindAllMarkers function and with the parameter “min.pct = 0.25, thresh.use = 0.25”.

### Single-cell pseudotime lineage trajectory reconstruction

Single-cell lineage reconstruction analysis was performed using Monocle (V2) packages according to the online tutorial (http://cole-trapnell-lab.github.io/monocle-release/docs/). The monocle object was constructed from Seurat object with newCellDataSet function, and Seurat determined variable genes were used as ordering genes to order cells in pseudotime along a trajectory. Dimension reduction was performed using DDRTree methods. To analyze differential gene expression between different cell branches, BEAM function was used and differentially expressed genes were identified with q-val < 1e-4. Branch-specific gene expression heatmap was plotted with plot_genes_branched_heatmap function and different gene set was calculated according to k-means clustering.

### Immunohistochemistry staining analysis

The immunofluorescence or enzyme horseradish peroxidase (HRP)-based immunohistochemistry analysis procedure was performed as we previously described (Ge et al., 2017, Liu et al., 2017). For immunofluorescence analysis, the paraffin-embedded skin tissues were firstly deparaffinized in xylene and further rehydrated in ethanol solutions. Antigen retrieval was performed in 0.01 M sodium citrate buffer at 96°C. Following a permeabilization procedure in 0.5 M Tris-HCI buffer supplemented with 0.5% TritonX-100 (Sorlabio, Beijing, China) for 10 min, the slides were then blocked with 3 % BSA and 10 % donkey serum (Boster, Wuhan, China) in 0.5 M Tris-HCI buffer for 30 min. The primary antibodies diluted in the blocking buffer were incubated with the slides at 4°C overnight. The next morning, the corresponding secondary antibodies were then added in the slides and incubated at 37°C for 1 h. DAPI was used to stain the nuclei and the pictures were taken using Nikon AR1 confocal system (Nikon, Tokyo, Japan). For HRP based immunohistochemistry analysis, following antigen retrieval, the samples were firstly incubated with 3% H_2_O_2_ for 10 min at room temperature to remove endogenous peroxidase activity. The primary antibodies were incubated 4 °C overnight, and the corresponding HRP-labeled secondary antibodies were added in the next morning for 40 min at room temperature. After that, peroxidase substrate DAB (Zsbio, Beijing, China) was used for chromogenic reaction with hematoxylin used to stain nuclei. The slides were finally mounted with neutral resins and pictures were captured with an Olympus BX51 microscope imaging system (Olympus, Tokyo, Japan). All the primary and secondary antibodies used in this study were listed in Supplementary Table 5.

### Data availability

The single-cell RNA sequencing data used in this research is deposited in NCBI GEO databases under accession number: GSE144351.

## Acknowledgments

This work was supported by the National Natural Science Foundation of China (31972556 and 31671554).

## Supplementary Figure Legends

**Supplementary Fig. 1 Single-cell dataset quality control and representative marker expression across all single cells.** (a) Morphology of fetus Cashmere goat at E60, E90 and E120 and its corresponding skin structure revealed by HE staining. (b) Quality matrices revealed by CellRanger of all datasets used in this study. (c) Comparison of the number of genes detected and the number of UMI detected in each cell. (d) Evaluating key cell type markers across all single cells in the tSNE plot.

**Supplementary Fig. 2 Gene function enrichment of differentially expressed genes during DC fate commitment.** (a) The network of enriched GO terms for different gene sets during DC fate commitment. Each dot represents one GO terms and different colors represent different gene sets. (b) Circos plot demonstrating the number of overlapped GO terms between different gene sets.

**Supplementary Fig. 3 Gene function enrichment of differentially expressed genes during DP cell fate commitment.** (a) The network of enriched GO terms for different gene sets during DP fate commitment. Each dot represents one GO terms and different colors represent different gene sets. (b) Circos plot demonstrating the number of overlapped GO terms between different gene sets. (c) Comparison of enriched GO terms between two different cell fate.

**Supplementary Fig. 4 Comparison of enriched GO terms during matrix cell fate commitment.** (a) Heatmap illustrating gene set specific enriched GO terms. (b) Circos plot demonstrating the number of overlapped GO terms between different gene sets.

**Supplementary Fig. 5 Validation of hair shaft/IRS marker expression and comparison of enriched GO terms.** (a) Immunohistochemistry analysis of hair shaft marker HOXC13 and IRS marker SOX9 expression in Cashmere goat skin. Scale bars = 50 μm. (b) The network of enriched GO terms for different gene sets during hair shaft and IRS fate commitment. Each dot represents one GO terms and different colors represent different gene sets. (c) Heatmap illustrating gene set specific enriched GO terms.

**Supplementary Fig. 6 Gene functional enrichment and representative gene expression along pseudtotime.** (a) The network of enriched GO terms for different gene sets during keratinocyte differentiation. Each dot represents one GO terms and different colors represent different gene sets. (b) Psedotime expression pattern of representative cell fate 1 and cell fate 2 signature genes.

## Supplementary Tables

**Supplementary table 1:** All cluster specific differentially expressed genes.

Supplementary table 2: DC signature genes for different gene sets.

Supplementary table 3: DC signature genes comparasion between goat and mice.

Supplementary table 4: IRS and hair shaft signature genes comparasion between goat and mice.

Supplementary table 5: All antibodies used in this study.

