## Supplemental Figures for "A single-cell transcriptome atlas during Cashmere goat hair follicle morphogenesis"

### Supplementary Figure 1

(a)

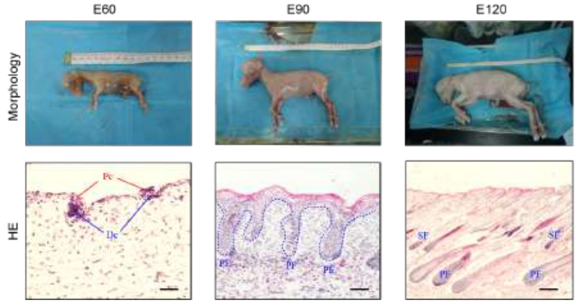


(b)

| Sample info. | E60 | E90 | E120 |
|---|---|---|---|
| Estimated Number of Cells | 7,000 | 7,000 | 7,000 |
| Valid Barcodes | 97.9% | 97.4% | 97.5% |
| Mean Reads per Cell | 83,588 | 218,817 | 81,753 |
| Median Genes per Cell | 1,822 | 1,202 | 2,289 |
| Total Genes Detected | 16,750 | 17,012 | 17,033 |
| Reads Mapped to Genome | 96.5% | 91.2% | 96.5% |

(c)

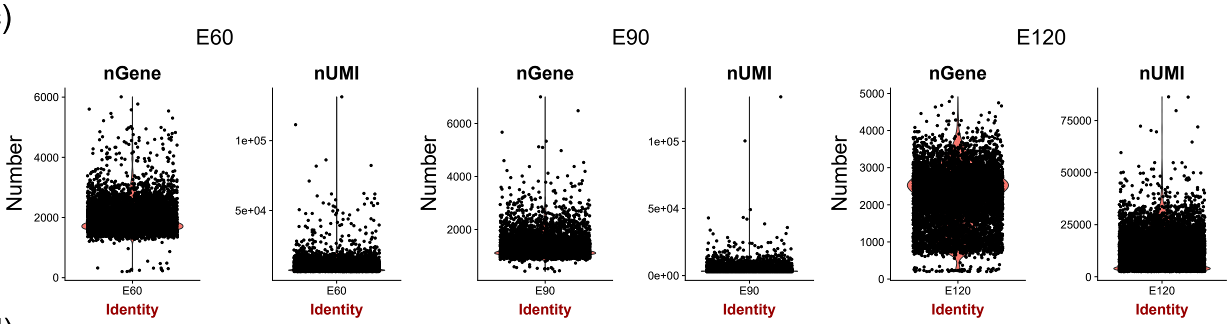


(d)

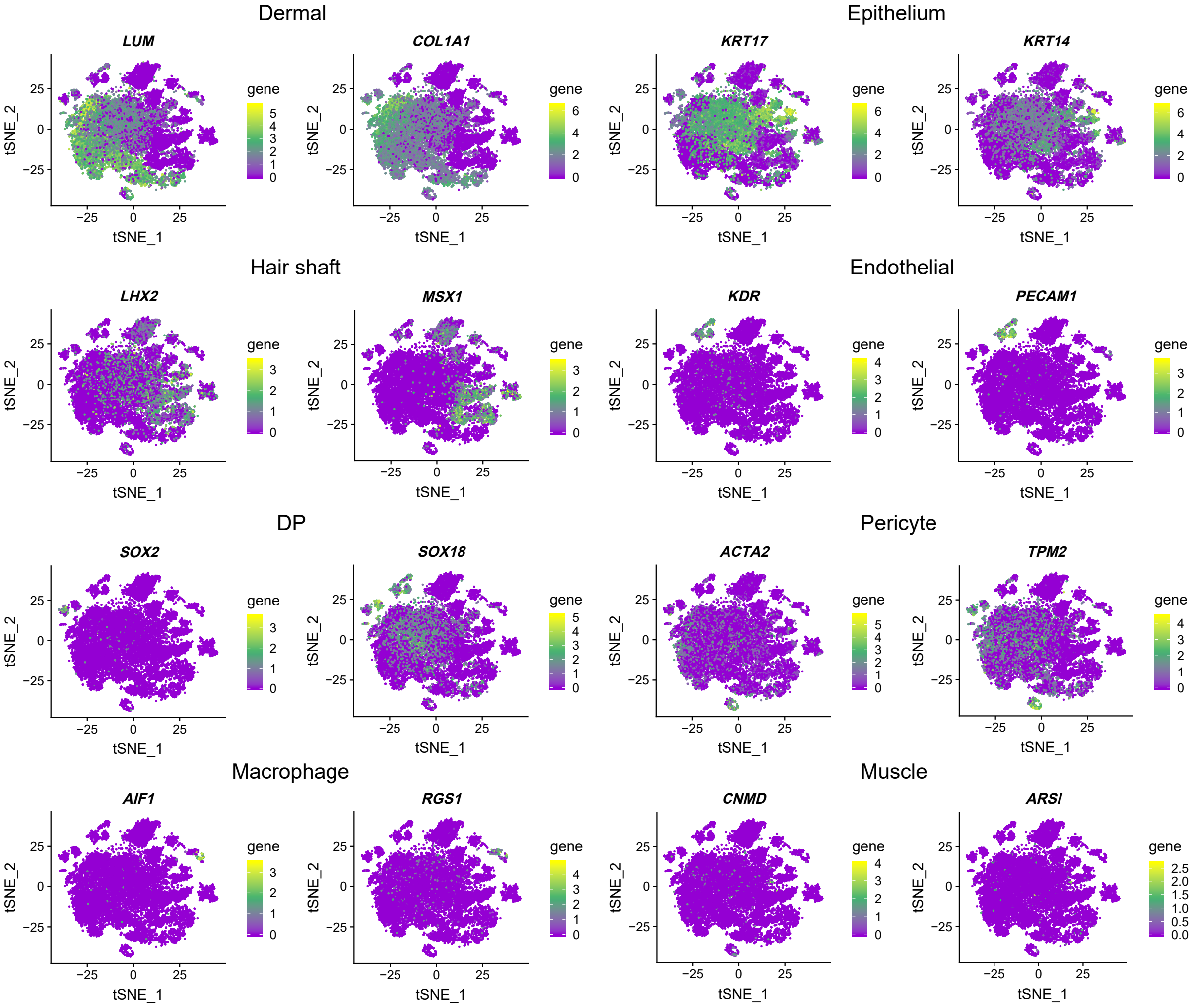

### Supplementary Figure 2

(a)

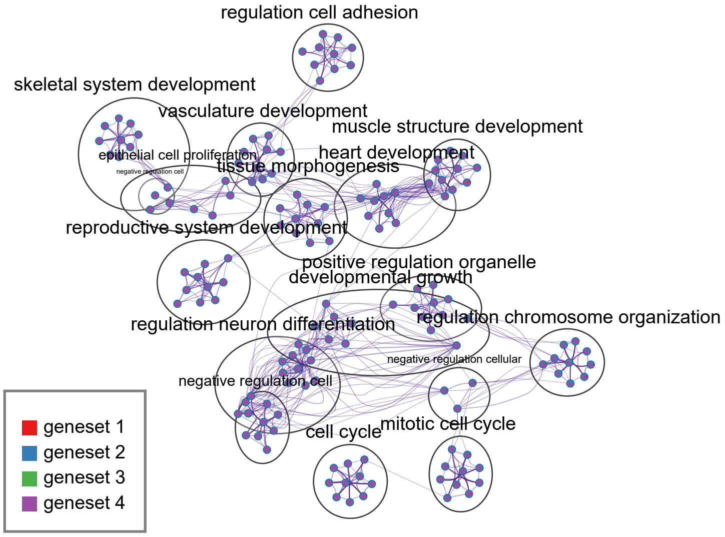


(b)

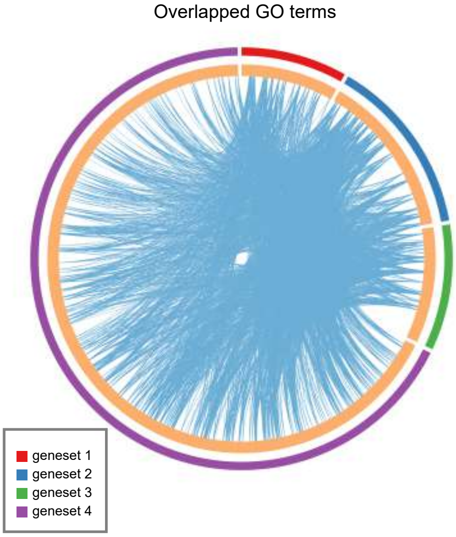

### Supplementary Figure 3

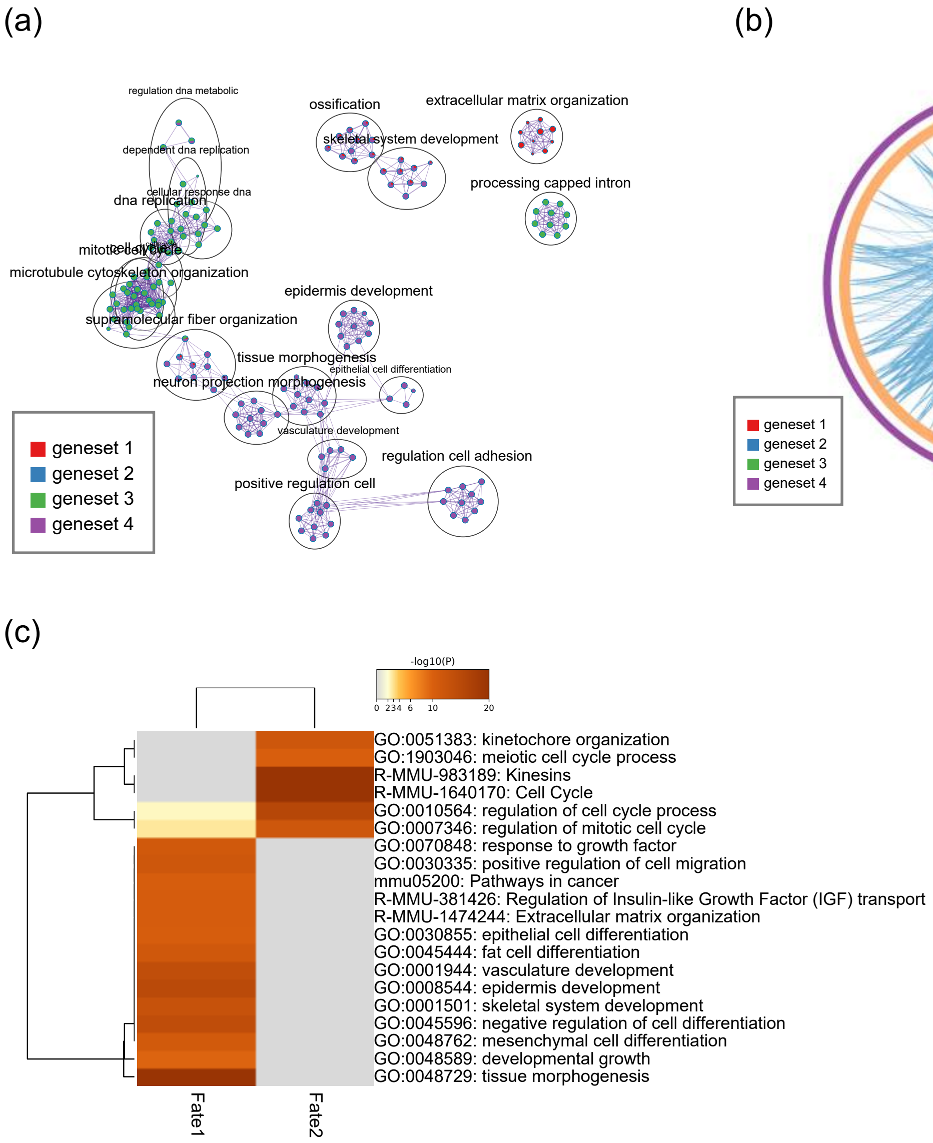

### Supplementary Figure 4

(a)

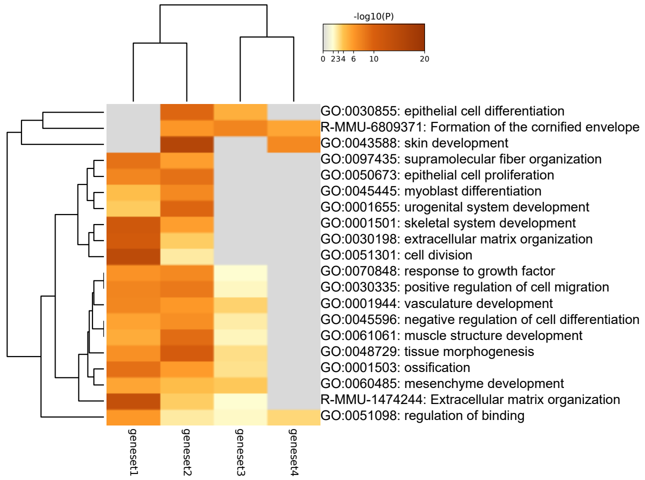


(b) Overlapped GO terms

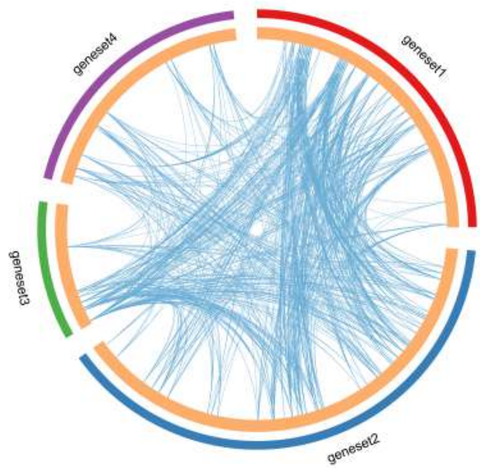

Supplementary Figure 5

(a)

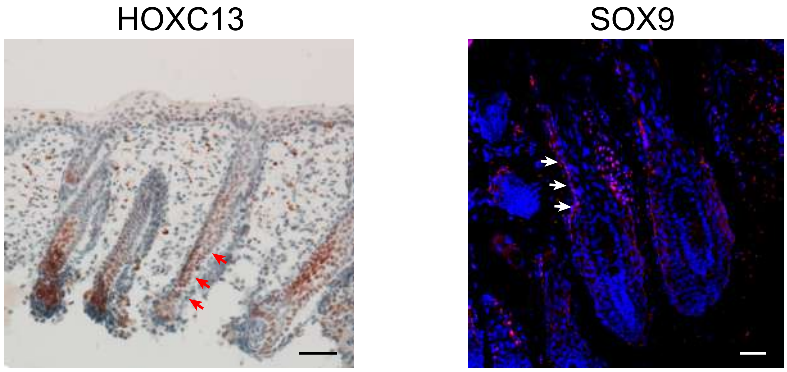


(b)

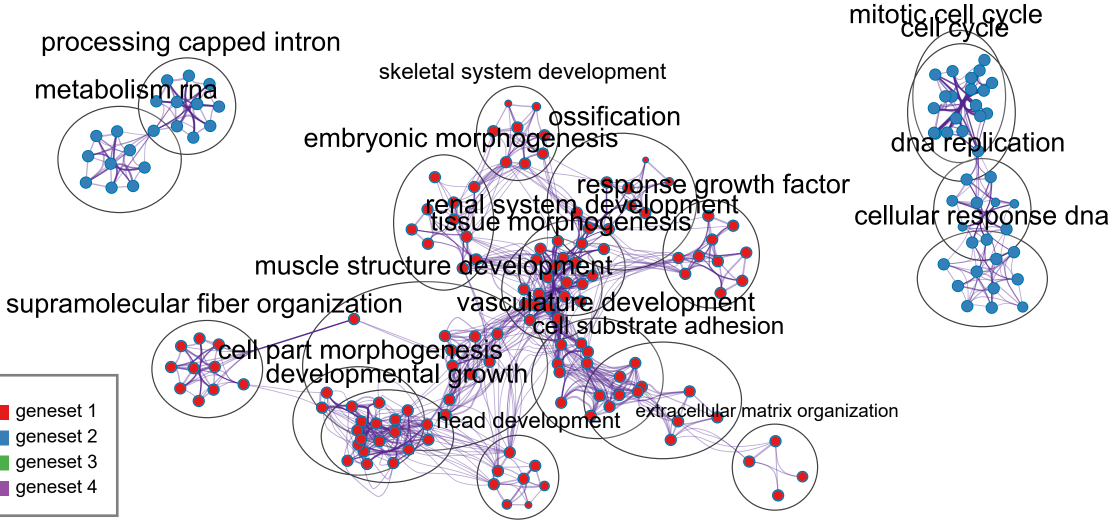


(c)

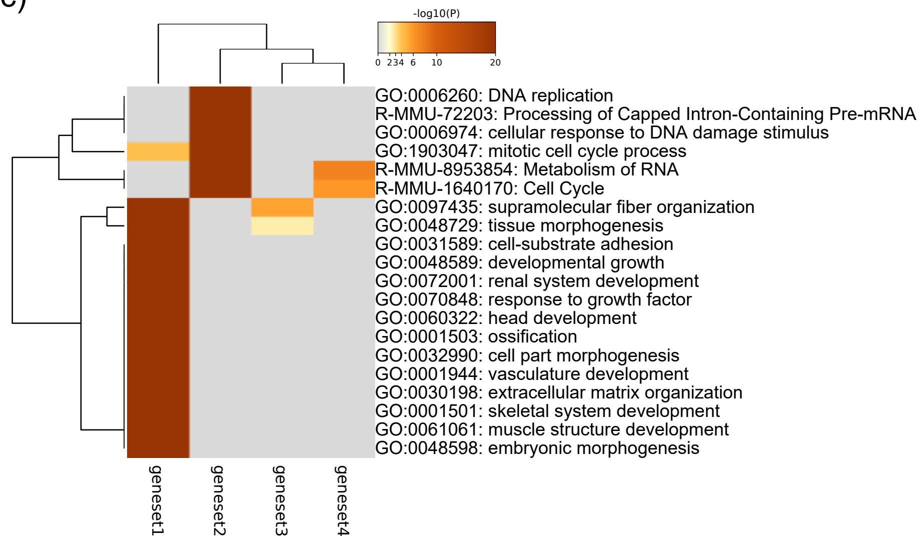

Supplementary Figure 6

(a)

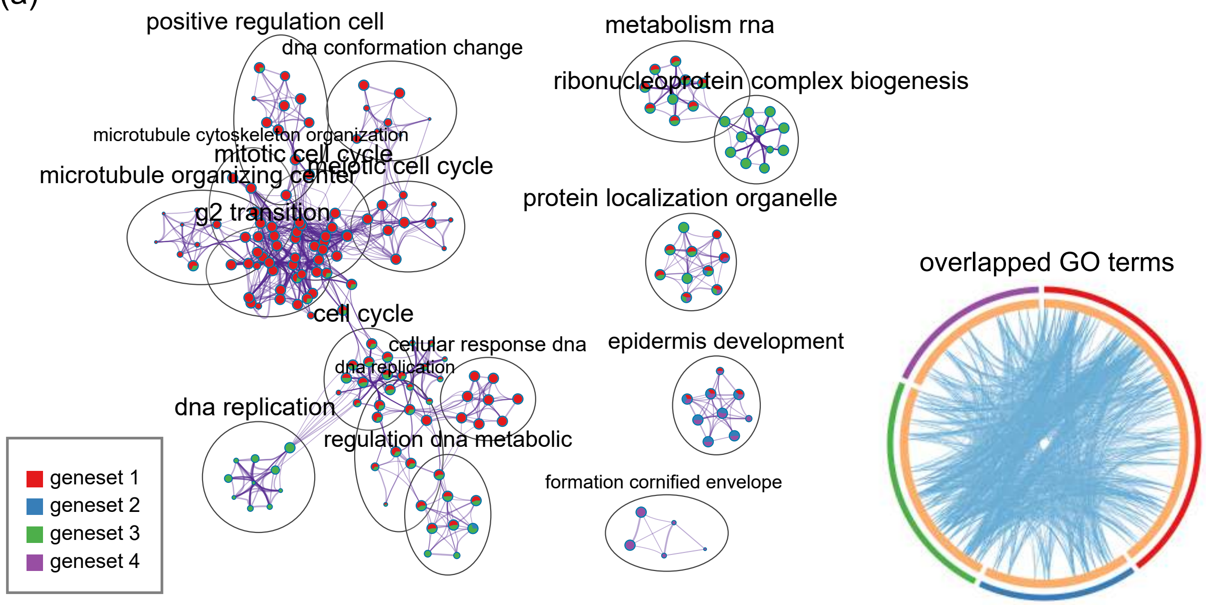


(b)

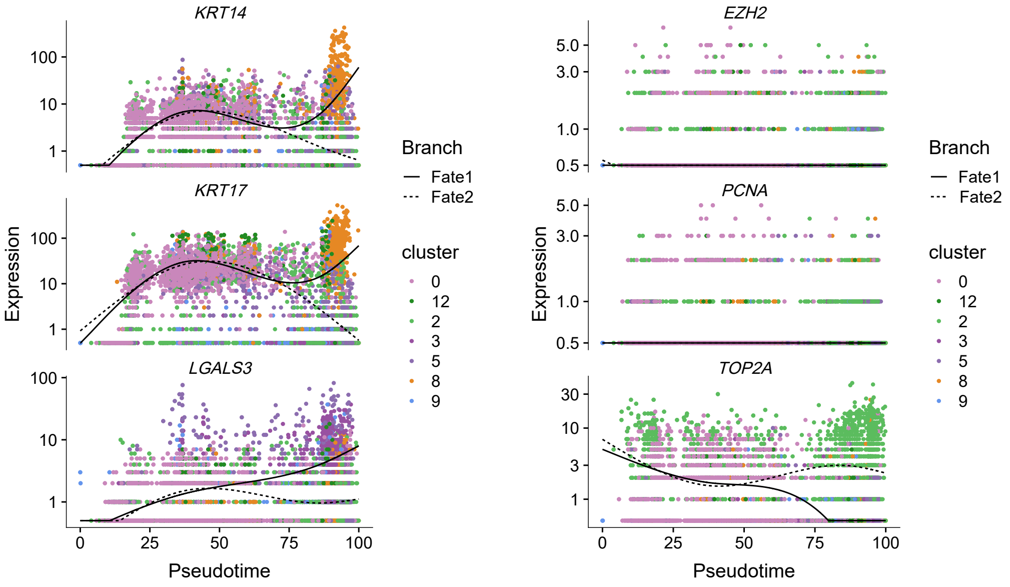
